# T cell priming is enhanced by maturation-dependent stiffening of the dendritic cell cortex

**DOI:** 10.1101/680132

**Authors:** Daniel Blumenthal, Lyndsay Avery, Vidhi Chandra, Janis K. Burkhardt

## Abstract

T cell activation by dendritic cells (DCs) involves forces exerted by the T cell actin cytoskeleton, which are opposed by the cortical cytoskeleton of the interacting APC. During an immune response, DCs undergo a maturation process that optimizes their ability to efficiently prime naïve T cells. Using atomic force microscopy, we find that during maturation, DC cortical stiffness increases via process that involves actin polymerization. Using stimulatory hydrogels and DCs expressing mutant cytoskeletal proteins, we find that increasing stiffness lowers the agonist dose needed for T cell activation. CD4^+^ T cells exhibit much more profound stiffness-dependency than CD8^+^ T cells. Finally, stiffness responses are most robust when T cells are stimulated with pMHC rather than anti-CD3ε, consistent with a mechanosensing mechanism involving receptor deformation. Taken together, our data reveal that maturation-associated cytoskeletal changes alter the biophysical properties of DCs, providing mechanical cues that costimulate T cell activation.

## INTRODUCTION

The initiation of an adaptive immune response requires priming of naïve T cells by professional antigen presenting cells (APCs). This process involves multiple receptor-ligand interactions, which occur in concert at a specialized cell-cell contact site called the immunological synapse (Dustin, 2014). Through these interactions, APCs transmit a highly orchestrated series of signals that induce T cell activation and direct differentiation of T cell populations (Friedl, den Boer, & Gunzer, 2005). While the biochemical aspects of this process have been the subject of many studies, the contribution of mechanical cues is only now being uncovered.

Following initial T cell receptor (TCR) engagement, T cells apply pushing and pulling forces on interacting APCs (Bashour et al., 2014; Hui, Balagopalan, Samelson, & Upadhyaya, 2015; Husson, Chemin, Bohineust, Hivroz, & Henry, 2011; Sawicka et al., 2017). These forces are essential for proper T cell activation (Li et al., 2010; Pryshchep, Zarnitsyna, Hong, Evavold, & Zhu, 2014). Moreover, force application is responsible, at least in part, for the ability of T cells to rapidly discriminate between agonist and antagonist antigens (Das et al., 2015; Liu, Chen, Evavold, & Zhu, 2014). While the mechanism by which force is translated into biochemical cues remains controversial (Das et al., 2015; Hong et al., 2015; Kim et al., 2009), there is evidence that early tyrosine phosphorylation events downstream of TCR engagement occur at sites where applied force is maximal (Bashour et al., 2014). Interestingly, the amount of force a T cell applies is directly affected by the stiffness of the stimulatory substrate (Husson et al., 2011; Sawicka et al., 2017). Thus, it appears that force application is mechanically coupled to the T cell’s ability to sense stiffness (mechanosensing). In other cell types, substrate stiffness has been shown to affect a variety of cell functions including differentiation, migration, growth and survival (Byfield, Reen, Shentu, Levitan, & Gooch, 2009; Discher, Janmey, & Wang, 2005; Engler, Sen, Sweeney, & Discher; Lo, Wang, Dembo, & Wang; Oakes et al., 2009; Pelham & Wang, 1997; Solon, Levental, Sengupta, Georges, & Janmey; Trappmann et al., 2012). Stiffness sensing in T cells has not been well studied, though there is some evidence that substrate stiffness affects both initial priming and effector functions (Basu et al., 2016; Judokusumo, Tabdanov, Kumari, Dustin, & Kam, 2012; O’Connor et al., 2012; Saitakis et al., 2017). Since the physiologically relevant substrate for T cell priming is the surface of the interacting APC, one might predict that changes in cortical stiffness of the APC will profoundly influence T cell priming. However, this prediction remains untested, and studies addressing the role of substrate stiffness in T cell priming did not take into consideration the physiological stiffness of APCs.

Dendritic cells (DCs) are the dominant APCs that prime T cells *in vivo* (Jung et al., 2002). One of the hallmarks of DC biology is the process of maturation. Immature DCs are sentinels of the immune system, specialized for immune surveillance and antigen processing (Mellman & Steinman, 2001). In response to infection or injury, inflammatory stimuli trigger signaling pathways that induce molecular reprogramming of the cell. The resulting mature DCs express high levels of surface ligands and cytokines needed for efficient T cell priming (Burns et al., 2004). The maturation process is tightly associated with remodeling of the DC actin cytoskeleton. This process underlies other maturation-associated changes such as downregulation of endocytosis and increased migratory behavior (Garrett et al., 2000; West, Prescott, Eskelinen, Ridley, & Watts, 2000). In addition, cytoskeletal remodeling has a direct impact on the ability of mature DCs to prime T cells (Al-Alwan, Rowden, Lee, & West, 2001; Comrie, Li, Boyle, & Burkhardt, 2015). Indeed, depolymerization of actin filaments perturbs the ability of mature peptide-pulsed DCs to activate T cells, indicating that actin plays an important role on the DC side of the immunological synapse. We hypothesized that maturation-associated changes in the actin cytoskeleton modulate the stiffness of the DC cortex, and promote T cell priming by providing physical resistance to the pushing and pulling forces exerted by the interacting T cell.

In this work, we aimed to better understand the relationship between DC cortical stiffness and T cell activation. We show that during maturation, DCs undergo a 2-3 fold increase in cortical stiffness, and that T cell activation is sensitive to stiffness over the same range. Moreover, we find that stiffness sensitivity is a general trait exhibited by most T cell populations. Since mechanosensing occurs lowers the threshold signal required for T cell activation, we conclude that stiffness serves as a novel biophysical costimulatory mechanism that functions in concert with canonical signaling cues to facilitate T cell priming.

## RESULTS

### Dendritic cell stiffness increases upon maturation

During maturation, DCs undergo a set of phenotypic changes that transform them into highly effective APCs (Mellman & Steinman, 2001). We hypothesized that as part of this maturation process, DCs might also modulate their cortical stiffness. To test this, we used atomic force microscopy (AFM) to directly measure cortical stiffness of immature and mature DCs. Murine bone marrow derived DCs (BMDCs) were prepared as described in Materials and Methods and cultured in the absence or presence of LPS to induce maturation. Cells were plated on Poly L-lysine (PLL) coated coverslips and allowed to spread for at least 4 hours prior to measurement of cortical stiffness by AFM micro-indentation. Because the population of LPS-treated cells was heterogeneous with respect to maturation markers, cells were labeled with fluorescent anti-CD86, and immature (CD86-negative) or mature (CD86 high) cells were selected for AFM measurements. As shown in Figure 1A, immature BMDCs were quite soft, with a mean Young’s modulus of 2.2±0.7 kPa. Mature BMDCs were almost two-fold stiffer, with a Young’s modulus of 3.5±1.0 kPa. Importantly, the stiffness of CD86-negative BMDCs within the LPS-treated population was the same as that of untreated, immature DCs. This demonstrates that the observed increase in stiffness is a property of DC maturation rather than an unrelated response to LPS treatment. Since BMDCs do not recapitulate all of the properties of classical, tissue resident DCs (Guilliams & Malissen, 2015; Lutz, Inaba, Schuler, & Romani, 2016; Na, Jung, Gu, & Seok, 2016), we verified our results by measuring the stiffness of *ex-vivo* DCs purified from spleens of untreated or LPS-injected mice. Results were very similar; the stiffness of immature splenic DCs was nearly identical to that of immature BMDCs, and LPS treatment resulted in an increase in stiffness of almost 3-fold (Figure 1B). These results demonstrate that stiffness modulation is a *bona fide* trait of DC maturation. Moreover, they establish that the biologically relevant range of DC stiffness lies between 2 and 8 kPa.

**Figure 1.**
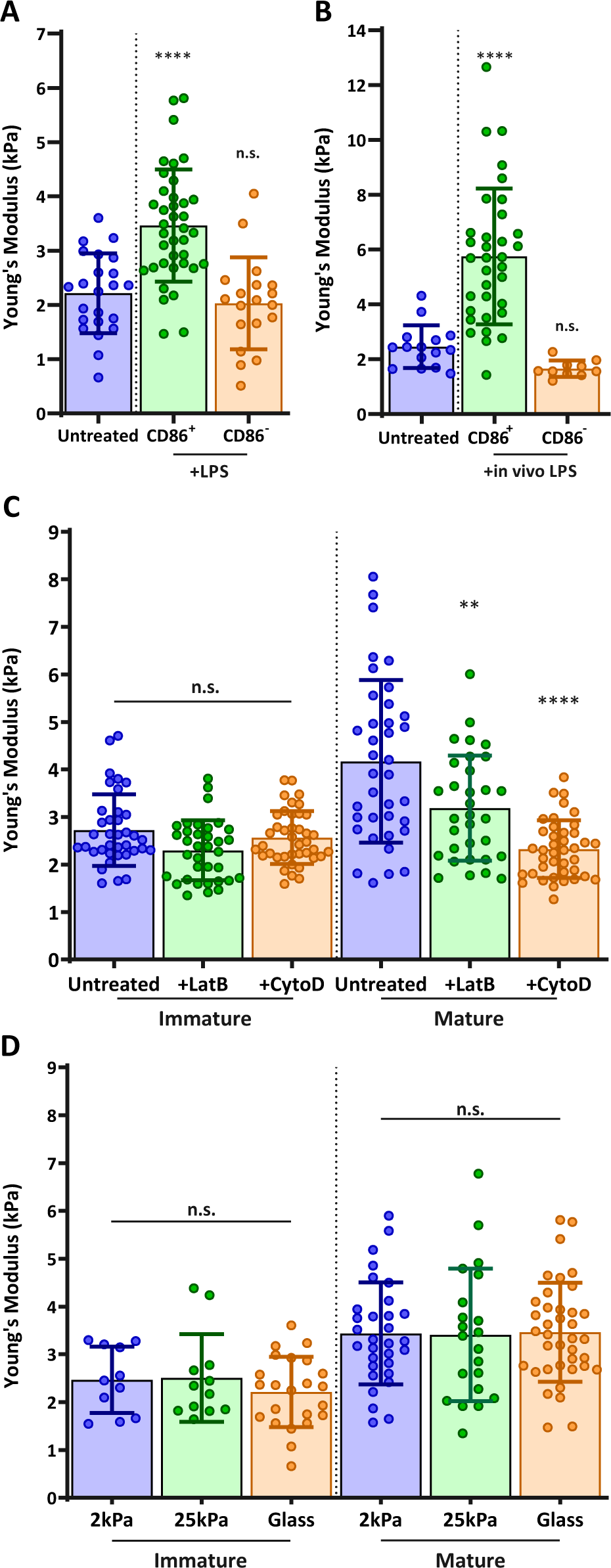
DC maturation induces an actin-dependent increase in cortical stiffness. (A) BMDCs were untreated or matured by treatment with LPS, and cortical stiffness was measured by AFM micro-indentation. Fluorescent anti-CD86 labeling was used to select immature (CD86-negative) or mature (CD86 high) cells. (B) E*x-vivo* DCs were purified from spleens of untreated mice or from mice injected with LPS 24 hours prior to harvesting the spleen, and analyzed as in A. (C) Immature or LPS matured BMDCs either left untreated or treated with 10µm of the actin de-polymerizing agents Latrunculin-B or Cytochalasin-D prior to AFM measurements. (D) Immature or LPS matured BMDCs were plated on substrates of different stiffness prior to AFM measurements. Data are pooled from 2-3 independent experiments. Each data point represents an average of two stiffness measurements at different locations around a single cell nucleus. Error bars denote standard deviation. n.s non-significant, **p<0.01, ****p<0.0001 calculated by an unpaired one-way ANOVA, post-hoc Tukey corrected test.

### The maturation-induced increase in stiffness is actin dependent and substrate independent

One well-known feature of DC maturation is remodeling of the actin cytoskeleton. This process involves changes in the activation status of Rho GTPases and downstream actin regulatory proteins, and is known to downregulate antigen uptake and increase cell motility (Garrett et al., 2000; West et al., 2000). To ask if changes in actin cytoarchitecture also result in increased cortical stiffness, we treated immature and mature BMDCs with the actin-depolymerizing agents Cytochalasin-D or Latrunculin-B. Neither drug affected the stiffness of immature BMDCs, indicating that the basal level of stiffness depends on factors other than the actin cytoskeleton (Figure 1C). In contrast, both drugs induced a significant decrease in the stiffness of mature DCs, with Cytochalasin reducing their stiffness to that of immature DCs. We conclude that the increased cortical stiffness observed upon DC maturation is another feature of actin cytoskeletal reprogramming.

Some cell types regulate their stiffness in response to the stiffness of their substrate (Byfield et al., 2009; Tee, Fu, Chen, & Janmey, 2011). To test whether DCs exhibit this behavior, immature and mature BMDCs were plated on PLL-coated substrates of different compliances (hydrogels of 2 or 25 kPa, or glass surfaces in the GPa range) and allowed to spread on the surface for at least 4 hours prior to AFM measurement. No apparent difference in cell spreading or morphology could be noted on the different PLL-coated hydrogels (data not shown). Hydrogel compliance was verified by measuring the elastic modulus of the surface in areas devoid of cells (Figure 1 - Figure Supplement 1). As shown in Figure 1D, substrate compliance had no effect on cortical stiffness of either immature or mature BMDCs. In control studies, we could readily detect substrate-dependent changes in stiffness of normal fibroblast cells (not shown). Thus, we conclude that DCs maintain a specific cortical stiffness, which is characteristic of their maturation state.

### DC cortical stiffness is primarily controlled by actin polymerization

We next sought to identify the molecular mechanisms controlling DC cortical stiffness. Several actin regulatory mechanisms are known to change during DC maturation. In particular, mature DCs upregulate the actin bundling protein fascin (Yamashiro, 2012), they show activation of myosin-dependent processes (van Helden et al., 2008), and they undergo changes in the activation and localization of Rho-GTPases, which in turn regulate actin polymerization via the Arp2/3 complex and formins (Burns, Thrasher, Blundell, Machesky, & Jones, 2001; Garrett et al., 2000; West et al., 2000). To ask how each of these pathways influences cortical stiffness, we used small molecule inhibitors and DCs from relevant knockout mice. Note that to facilitate comparison between experiments, control immature and mature DCs were tested in each experiment, and results were normalized based on values for mature DCs. First, we tested the role of fascin, which is known to generate very stiff actin bundles *in vitro* (Demoulin, Carlier, Bibette, & Baudry, 2014). Surprisingly, the stiffness of BMDCs from fascin^-/-^ mice was indistinguishable from that of WT BMDCs both before and after LPS-induced maturation (Figure 2A). Next, we tested the contribution of myosin contractility, which is known to control stiffness and membrane tension in other cell types (Salbreux, Charras, & Paluch, 2012). As shown in Figure 2B, treating mature BMDCs with the myosin II inhibitor blebbistatin reduced stiffness by a small, albeit statistically significant amount. Similar results were obtained with the Rho-kinase (ROCK) inhibitor Y27623, which indirectly inhibits myosin function.

**Figure 2.**
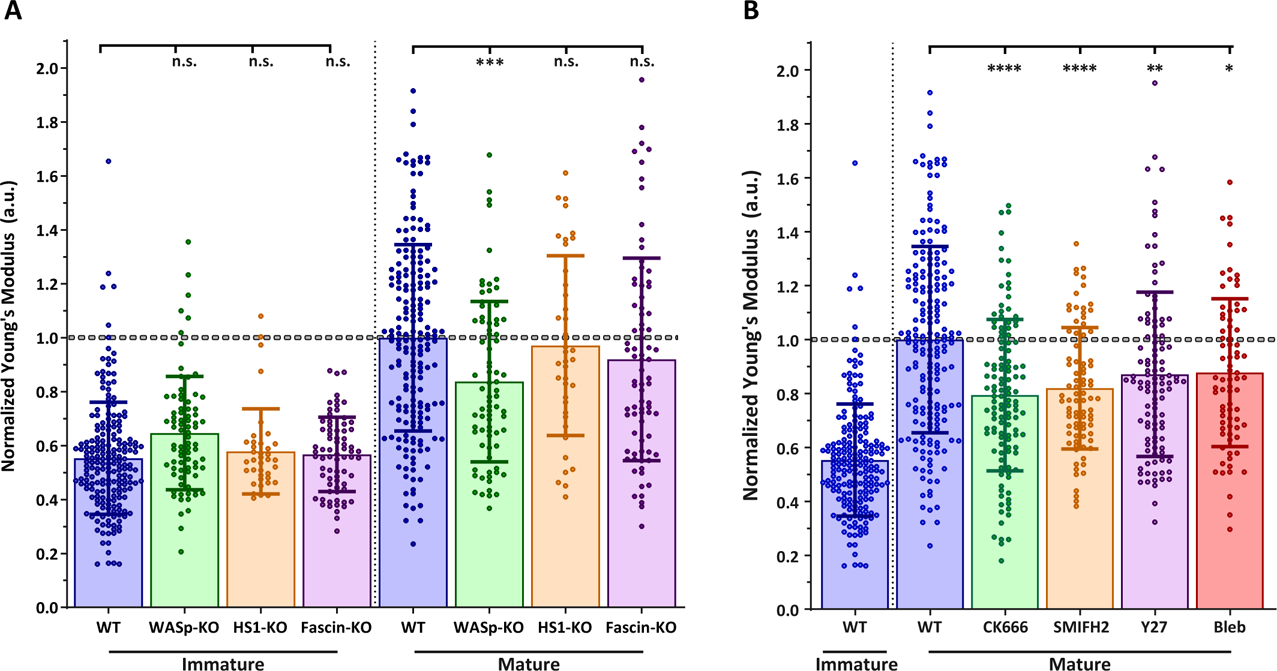
Effects of actin regulatory proteins on DC cortical stiffness. Murine BMDCs from WT mice or mice lacking key actin-associated proteins were untreated or matured by treatment with LPS, and cortical stiffness was measured by AFM micro-indentation. To facilitate comparison between experiments, results were normalized to values of mature WT BMDCs in each experiment. (A) Cortical stiffness of BMDCs from mice lacking important actin modulating proteins. (B) Cortical stiffness of WT BMDCs treated with cytoskeletal inhibitors. CK666 (100 µM) was used to inhibit branched actin polymerization by Arp2/3. SMIFH2 (10 µm) was used to inhibit linear actin polymerization by formins. Acto-myosin contractility was inhibited directly with Blebbstatin (50 µM) or indirectly with the Rho-kinase inhibitor Y27623 (25 µM). All drugs had no effect on immature BMDCs (data not shown). Data points for untreated WT BMDCs (both immature and mature) were pooled from all experiments as a reference. The dashed line represents the average stiffness of untreated mature WT BMDCs from all experiments. Data are pooled from 2-3 independent experiments. Each data point represents an average of two stiffness measurements at different locations around a single cell nucleus. Error bars denote standard deviation. n.s non-significant, *p<0.05, **p<0.01, ***p<0.001, ****p<0.0001 calculated by an unpaired one-way ANOVA, with post-hoc Tukey correction.

We next considered the possibility that cortical stiffness is modulated by actin polymerization. Broadly speaking, actin polymerization is induced by two sets of proteins: formins generate linear actin filaments, while activators of the Arp2/3 complex produce branched actin structures. Treatment of DCs with the pan-formin inhibitor SMIFH2 significantly reduced the cortical stiffness of mature DCs (Figure 2B). A similar reduction was observed after inhibition of Arp2/3-mediated branched actin polymerization by CK666. DCs express multiple activators of Arp2/3 complex, of which two have been implicated in maturation-associated changes in actin architecture: Hematopoietic Lineage Cell-Specific Protein 1 (HS1), the hematopoietic homologue of cortactin (Huang et al., 2011), and WASp, the protein defective in Wiskott-Aldrich syndrome (Bouma, Burns, & Thrasher, 2007; Bouma et al., 2011; Calle, Chou, Thrasher, & Jones, 2004). To individually assess the role of these two proteins, we used BMDCs cultured from HS1 and WASp knockout mice. As shown in Figure 2A, loss of HS1 had no impact on cortical stiffness of either immature or mature BMDCs. In contrast, mature WASp knockout BMDCs were significantly less stiff than WT controls. This difference mirrors that seen after inhibition of Arp2/3 complex by CK666, suggesting that WASp is the primary activator of Arp2/3 complex-dependent changes in cortical stiffness. The defect in WASp knockout DCs was observed only after maturation; immature WASp knockout DCs did not differ in stiffness from WT controls. This is consistent with our finding that the stiffness of immature DCs is unaffected by actin depolymerizing agents. Taken together, these results show that activation of formin and WASp-dependent actin polymerization pathways, and to a lesser extent increased myosin contractility, all contribute to the increased cortical stiffness of mature DCs.

### Stiffness lowers the threshold for activation of CD4^+^ T cells, but not CD8^+^ T cells

Our findings raise the possibility that changes in DC cortical stiffness, like other maturation-induced changes, enhance the ability of these cells to prime a T cell response. Previous studies have shown that T cells are sensitive to the stiffness of stimulatory surfaces (Judokusumo et al., 2012; O’Connor et al., 2012), and that that the TCR serves as the mechanosensory (Judokusumo et al., 2012). However, the results are conflicting, and these earlier studies were not performed within the physiological stiffness range that we have defined for DCs. Thus, we tested T cell responses on hydrogels with a stiffness range spanning that of immature and mature DCs (2 – 25 kPa). We verified the compliance of the hydrogel surfaces by measuring the elastic modulus of the surfaces directly by AFM. Hydrogel stiffnesses were found to be similar to those reported by the manufacturer (Figure 3 - Figure Supplement 1). Surfaces were coated with varying doses of peptide-loaded major histocompatibility complex (pMHC) molecules, together with a constant dose of anti-CD28. Surfaces were coated with H-2K^b^ class I MHC loaded with the N4 (SIINFEKL) peptide (pMHC-I), or I-A^b^ class II MHC loaded with the OVA_329-337_ (AAHAEINEA) peptide (pMHC-II), to stimulate OT-I CD8^+^ or OT-II CD4^+^ T cells, respectively. Plastic surfaces, which are commonly used for stimulation with surface-bound ligands, were included as a familiar reference point. To test the effects of substrate stiffness on early T cell activation, we measured surface expression of the activation markers CD25 and CD69, as well as the production of IL-2, all at 24 hours post stimulation. As shown in Figure 3A-C, CD4^+^ T cells showed a profound stiffness-dependent response at 24 hours across all measures. This response was most clearly seen for upregulation of CD25 and CD69, where increasing substrate stiffness enhanced the CD4+ T cell response in a graded manner. As expected, for any given substrate stiffness T cell activation increased with increasing peptide dose. However, comparison between stimulatory surfaces revealed that increasing substrate stiffness lowered the pMHC-II dose required to obtain the same level of activation. Over the stiffness range associated with DC maturation (2-8kPa), the dose of TCR signal needed to induce surface marker upregulation was shifted by 1-2 logs. Analysis of IL-2 production revealed a similar effect, although the stiffness sensitivity was more bi-modal. IL-2 production increased almost 3-fold when CD4+ T cells were stimulated on surfaces of 8kPa, as opposed to 2kPa, for the same antigen dose.

**Figure 3.**
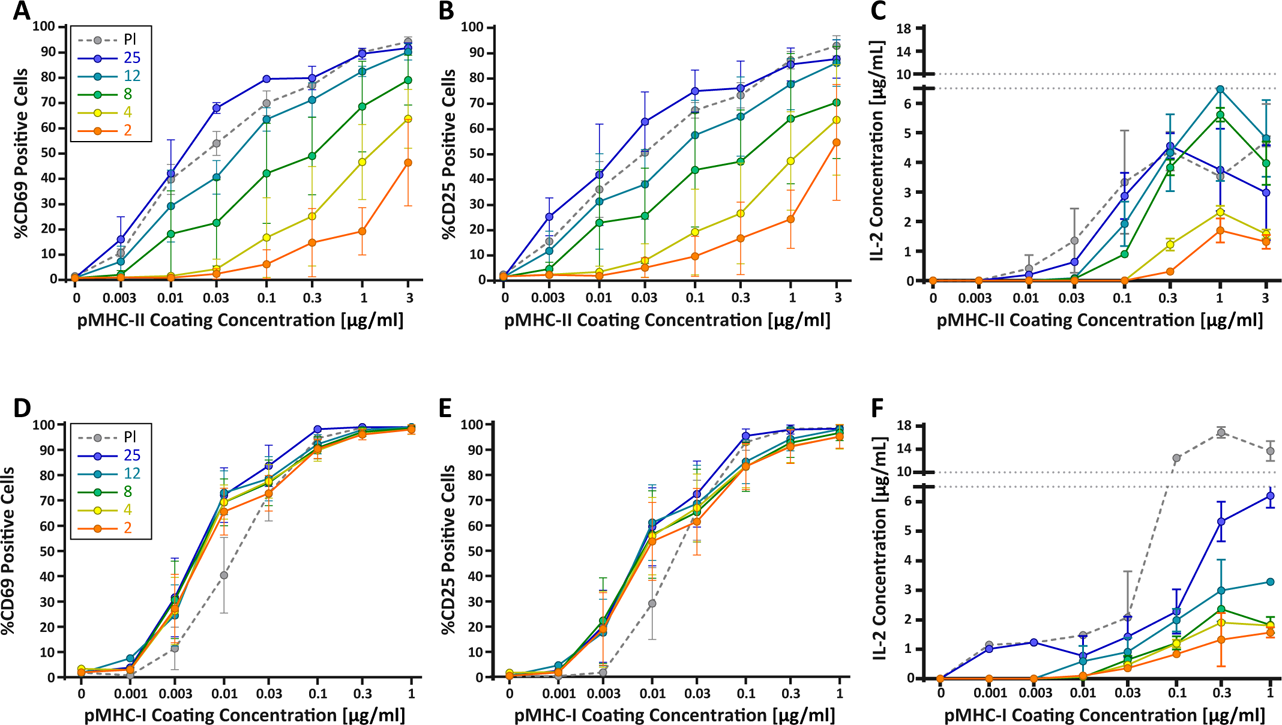
CD4^+^ and CD8^+^ T cells demonstrate vastly different stiffness responses. Murine T cells were purified from lymph nodes and spleen and activated on stimulatory acrylamide hydrogel surfaces with a stiffness range of 2 – 25 kPa (and plastic). Stimulatory surfaces were coated with the indicated concentrations of pMHC-I or pMHC-II together with 2 µg/ml anti-CD28 to stimulate OT-I CD8^+^ or OTI-II CD4^+^ T cells, respectively. (A,B,D,E) Cells were harvested 24 hours post activation and expression of early activation markers was measured by flow cytometry. Data represent averages +/-SEM of percent positive cells from N=3 independent experiments. (A,B) CD4^+^ T cells show profound stiffness dependent expression of both markers. (C,F) Cell supernatants were collected at 24 hours and IL-2 expression was analyzed by ELISA. Data represent means +/-StDev from 3-6 replicate samples from one representative experiment, N=2 experiments.

Strikingly, the robust stiffness response we observed in CD4^+^ T cells was not recapitulated for CD8^+^ T cells, especially when T cell activation was assessed based on surface marker upregulation (Figure 3D,E). Analysis of IL-2 production did reveal stiffness-enhanced activation of naïve CD8^+^ T cells, but this was largely restricted to substrates outside the stiffness range defined for DCs (12kPa, 25 kPa and plastic).

To determine if the stiffness dependency of CD4^+^ T cells seen at early times after TCR engagement is maintained at later times, we measured T cell proliferation based on CFSE dilution at 72 hours post stimulation. Similar to what was observed for early activation markers, increasing substrate stiffness produced graded increases in CD4^+^ T cell proliferation, and the threshold dose required to induce robust proliferation shifted as a function of substrate stiffness (Figure 4A,B). This effect was particularly evident at low doses of pMHC-II (0.1-1ug/ml). Interestingly, although soft hydrogels (2-4 kPa) elicited only very low levels of CD4^+^ T cell proliferation, these substrates did induce upregulation of CD25 in a high percentage of CD4^+^ T cells, even in the undivided populations (Figure 4C,D). This indicates that an activating signal was received, but was insufficient to drive proliferation.

**Figure 4.**
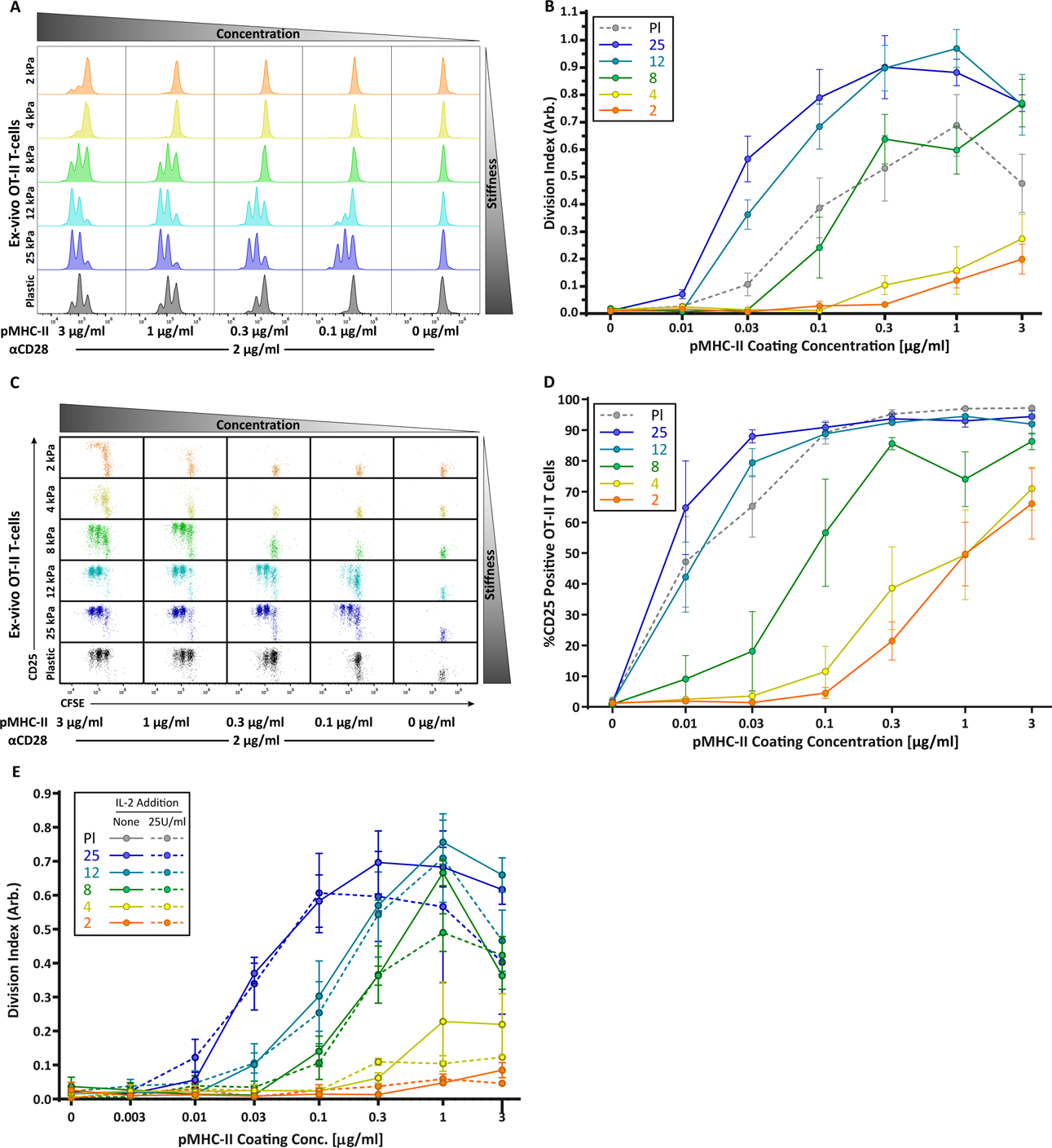
CD4 T cell proliferation is stiffness dependent and IL-2 independent. OT-II CD4^+^ T cells were purified from lymph nodes and spleen and activated on stimulatory acrylamide hydrogel surfaces with a stiffness range of 2 – 25 kPa (and plastic). Stimulatory surfaces were coated with the indicated concentrations of pMHC-II together with 2 µg/ml anti-CD28. (A,B) Proliferation of CD4^+^ T cells was measured by CFSE dilution at 72 hours post activation, showing profound stiffness-dependent proliferation. (A) Representative CFSE dilution matrix from a single experiment. (B) Average division index from 5 independent experiments. (C) Representative plots of CD25 expression as a function of CFSE dilution at 72 hours from a single experiment shows that upregulation of CD25 on T cell membrane precedes proliferation. (D) Average percent of T cells expressing CD25 from 5 independent experiments. (E) Division index of CD4^+^ T cells activated with or without addition of 25 U/ml of exogenous IL-2. Data in B, D, and E represent averages +/-SEM from at least 3 independent experiments.

Since the threshold stimuli (stiffness and dose of pMHC-II) required to induce significant IL-2 production and proliferation were very similar (Figures 3C and 4B), we reasoned that the threshold for proliferation might be driven by IL-2 availability. To test this, OT-II CD4^+^ T cells were stimulated on hydrogels with or without addition of 25 U/ml exogenous IL-2. Interestingly, addition of IL-2 did not rescue the proliferation of T cells stimulated on soft surfaces (Figure 4E). Thus, we conclude that in addition to influencing the signaling threshold for IL-2 production, substrate stiffness also affects other IL-2 independent events needed for efficient T cell proliferation.

Although early activation events in CD8^+^ T cells did not exhibit stiffness sensitivity, we reasoned that proliferative responses might behave differently. As shown in (Figure 5A,B), CD8^+^ T cells showed a mild stiffness-dependent proliferation response. The concentration of stimulatory ligand needed to induce at least one round of division was similar across the entire stiffness range Over successive rounds, increased stiffness did enhance the extent of proliferation, but the differences were relatively small (Figure 5B). Interestingly, analysis of CFSE dilution as a function of CD25 expression reveals evidence that CD8^+^ T cells exhibit a binary stiffness response (Figure 5 C,D); at low doses of peptide ligand, cells stimulated on very stiff substrates (25kPa or plastic) survived and proliferated, whereas cells stimulated on softer substrates were mostly lost (Figure 5C). From the standpoint of T cell priming, the significance of this observation is unclear, since these stiffnesses are well outside the biologically relevant range we measured for DCs.

**Figure 5.**
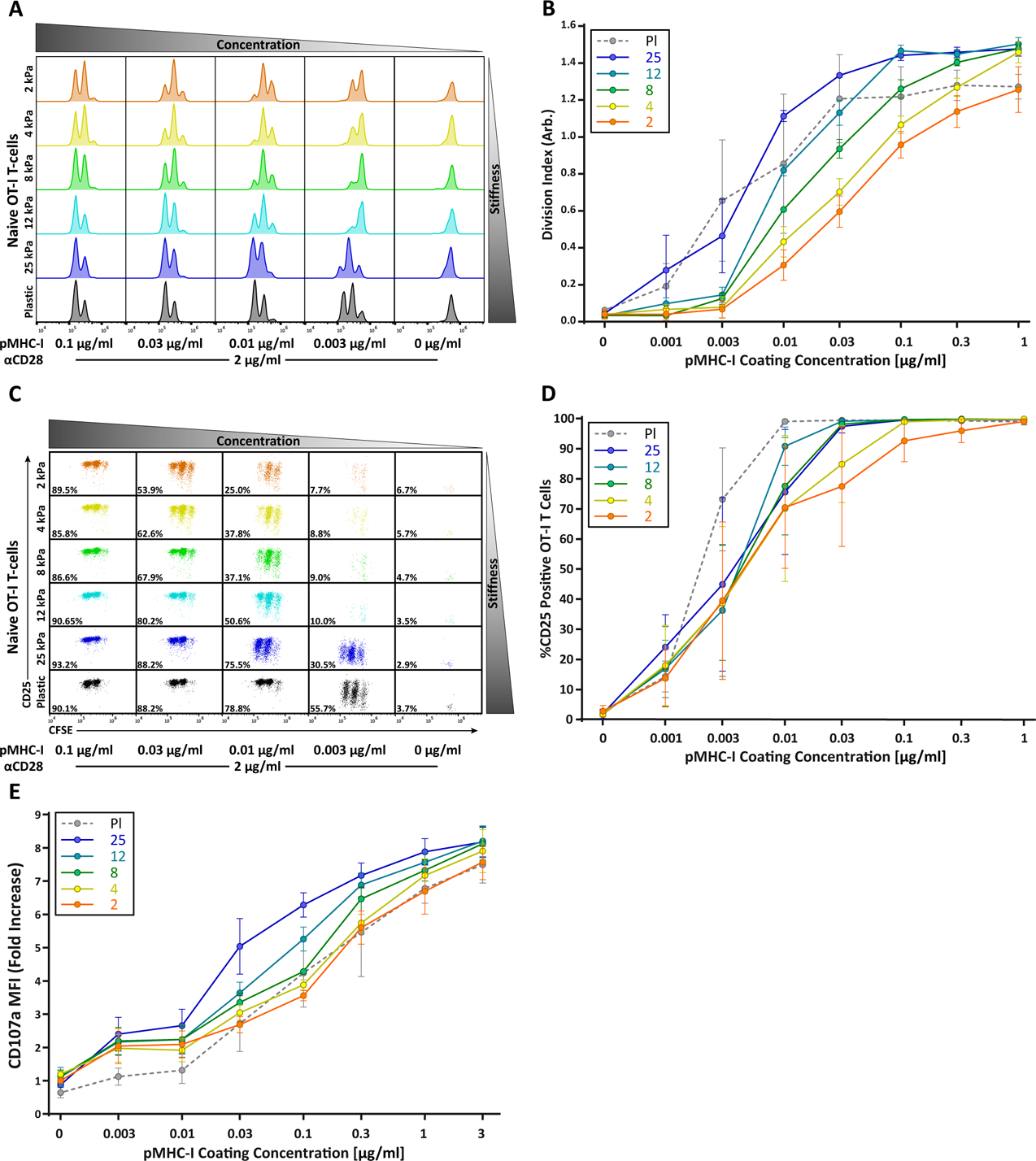
CD8^+^ T cell proliferation and degranulation show moderate stiffness dependency. Naïve OT-I CD8^+^ T cells were purified from lymph nodes and spleen and activated on stimulatory acrylamide hydrogel surfaces with a stiffness range of 2 – 25 kPa (and plastic). Stimulatory surfaces were coated with the indicated concentrations of pMHC-I together with 2 µg/ml anti-CD28 (A,B) Proliferation of CD8^+^ T cells was measured by CFSE dilution at 72 hours post activation, showing only moderate stiffness-dependent proliferation. (A) Representative CFSE dilution matrix from a single experiment. Note that the threshold pMHC-I concentration needed to induce proliferation is very similar between the different stiffness surfaces. (B) Average division index from 3 independent experiments. (C) representative plot of CD25 expression as a function of CFSE dilution at 72 hours from a single experiment shows a binary stiffness response. Note that with low doses of pMHC-I, only cells stimulated on very stiff substrates (25kPa or plastic) survived and proliferated. Percent of live cells is given for each condition. (D) Average percent of T cells expressing CD25 from 3 independent experiments. Levels of CD25 membrane expression are very similar between the different substrates, probably reflecting the fact that only T cells that upregulate CD25 survive. (E) Cytotoxic CD8^+^ T cells on days 8 or 9 of culture were restimulated on hydrogel surfaces with a range of pMHC-I concentrations, and degranulation was measured based on surface exposure of CD107a (N=3). Data in B, D, and E represent averages +/-SEM from at least 3 independent experiments.

Taken together, our findings point to a mechanism in which stiffer substrates have a sensitizing effect on CD4^+^ T cells, similar to that of classical co-stimulatory molecules such as CD28 (Harding, McArthur, Gross, Raulet, & Allison, 1992). When considered in this way, the relative lack of stiffness responses in CD8^+^ T cells fits with the fact that CD8^+^ T cells are much less dependent on costimulatory signals (McAdam, Schweitzer, & Sharpe, 1998).

### Degranulation of cytotoxic T cells shows mild stiffness sensitivity

Whereas naïve T cells are activated by DCs, effector T cells interact with many cell types. In particular, cytotoxic CD8^+^ T cells (CTLs) must respond to a variety of possible target cells, which may differ widely with respect to stiffness. We therefore reasoned that CTL effectors might be stiffness independent. To test this, *ex vivo* OTI CD8^+^ T cells were activated on plastic surfaces and grown in the presence of exogenous IL-2 to produce mature CTLs. To induce and detect release of cytolytic granules, CTLs were re-stimulated on hydrogels coated with a range of pMHC-I concentrations in the presence of fluorescent anti-CD107a antibody and analyzed by flow cytometry. Degranulation showed only a mild stiffness dependency (Figure 5E), and stiffness tended to affect the magnitude of degranulation rather than whether or not a degranulation response was triggered. Interestingly, changes in substrate stiffness within the range of defined for DCs (2-8kPa) had little or no impact on degranulation. Increased degranulation was only seen on stiffer surfaces (12 and 25 kPa). This could have functional consequences for effector function *in vivo*, since target cells in inflamed tissues can reach this stiffness range.

### Engagement the TCR complex by pMHC elicits the most prominent stiffness response

Many current models for TCR mechanosensing are founded on the notion of TCR deformation following engagement of cognate pMHC (Ma, Janmey, & Finkel, 2008). According to this concept, forces applied by the T cell on the TCR-pMHC bond result in conformational changes within TCRαβ (primarily the β chain), which are transmitted to intracellular components of the TCR/CD3 complex, leading to the initiation of downstream signaling (Das et al., 2015; Lee et al., 2015; Swamy et al., 2016). Given this, we wondered whether engaging the TCR complex through the CD3ε chain, as compared to direct TCRαβ engagement, would differentially affect T cell mechanosensing. To test this, CD4^+^ OT-II T cells were stimulated on hydrogels coated with anti-CD3ε antibodies, anti-TCRβ antibodies, or pMHC-II monomers. T cell activation was measured at 72 hours based on proliferation (CFSE dilution) and expression of CD25 (Figure 6). All three ligands induced a stiffness-dependent response for both proliferation and CD25 expression, but the responses differed in significant ways. In general, stimulating T cells with pMHC-II resulted in the strongest responses on the hydrogel surfaces; only on plastic surfaces did anti-CD3ε stimulation yield a similarly strong response (Figure 6A,C). (Figure 6A). Importantly, on surfaces with a stiffness similar to that of mature DCs (8kPa), stimulation with pMHC resulted in robust proliferation, while stimulation with anti-CD3ε yielded a minimal response. On softer substrates in the 2-4kPa range, pMHC elicited some proliferation, whereas stimulation with anti-CD3 did not. Interestingly, stimulation with anti-TCRβ resulted in a mixed response. At high doses of antigen, stimulation with anti-TCRβ yielded clear proliferative responses on substrates within the biologically-relevant stiffness range. Analysis of CD25 expression patterns revealed a similar trend (Figure 6D-F); on soft surfaces, pMHC yielded the strongest response, and anti-TCRβ was more effective than anti-CD3ε. Taken together, these results indicate that T cells sense substrate stiffness best through direct engagement of TCRαβ.

**Figure 6.**
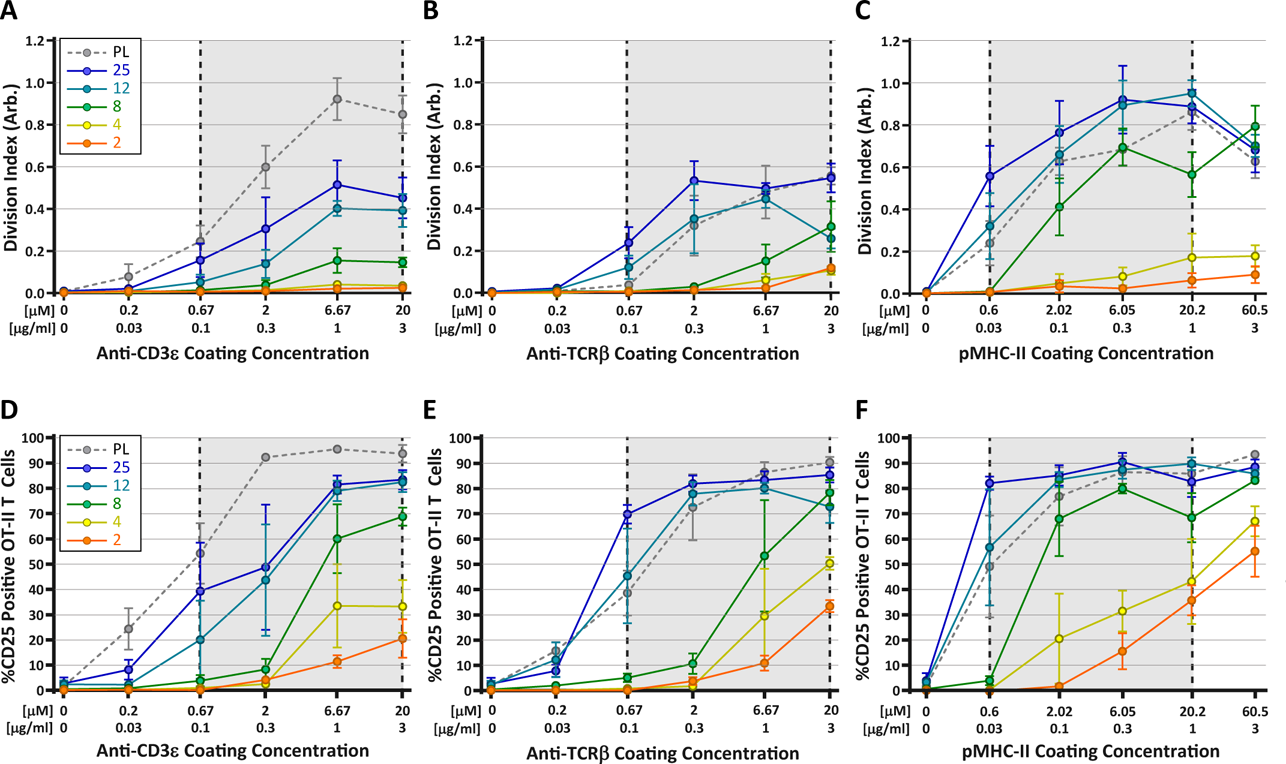
Stimulation of TCR with pMHC leads to the strongest stiffness-dependent response. OT-II CD4^+^ T cells were stimulated on acrylamide hydrogels of different stiffnesses coated with the indicated range of stimulatory ligands. Proliferation and membrane expression of CD25 were measured 72 hours post activation by flow cytometry. (A,D) Stimulation with anti-CD3ε antibody. (B,E) Stimulation with anti-TCRβ antibody. (C,F) Stimulation with pMHC-II. Plots show the average division index or CD25 expression from 3 independent experiment. Gray areas denote a similar range of stimulatory ligand molar concentrations to aid comparison.

### The increased stiffness of mature DCs enhances their ability to prime T cells

Our hydrogel assays show that T cell activation is enhanced by changes in stiffness over the range observed for DC maturation, consistent with the idea that modulation of cortical stiffness is a biophysical mechanism by which DCs control T cell activation. To test this directly, we sought conditions under which we could manipulate the stiffness of mature DCs. We took advantage of our finding that mature WASp knockout BMDCs are approximately 20% softer than WT controls (Figure 2A; data are presented as absolute values in Figure 7A). WT and WASp^-/-^ BMDCs were pulsed with increasing concentrations of OVA_323-339_ peptide and co-cultured with OT-II CD4^+^ T cells. T cell proliferation was measured by CFSE dilution. As shown in Figure 7B, WASp^-/-^ BMDCs did not prime T cells as efficiently as WT BMDCs at low OVA concentrations. Higher concentrations of OVA rescued this defect, showing that loss of WASp shifts the dose of peptide needed rather than affecting T cell priming *per se,* in keeping with the view that DC stiffness provides a costimulatory signal. We next attempted to test T cell priming activity of DCs that are stiffer than WT cells. We tested several genetic manipulations, most of which did not significantly increase the cortical stiffness of mature BMDCs. We did find that overexpression of a constitutively active form of WASp (I294T, CA-WASp (Beel et al., 2009)) increased cortical stiffness of mature BMDCs by approximately 20% relative to WT cells (Figure 7A), but these BMDCs failed to prime T cells more efficiently (Figure 7C). Expression of CA-WASp only enhances BMDC stiffness to approximately 5kPa, and based on our hydrogel studies, this increase is unlikely to be sufficient to enhance T cell activation. It seems likely that conditions that stiffen DCs to 10kPa or more would further enhance T cell responses, but we were unable to test this directly, and it is not clear whether this happens in vivo. Nonetheless, the studies using WASp-/-DCs show that changes in DC stiffness within the range observed during maturation has a significant impact on their ability to efficiently prime a T cell response.

**Figure 7.**
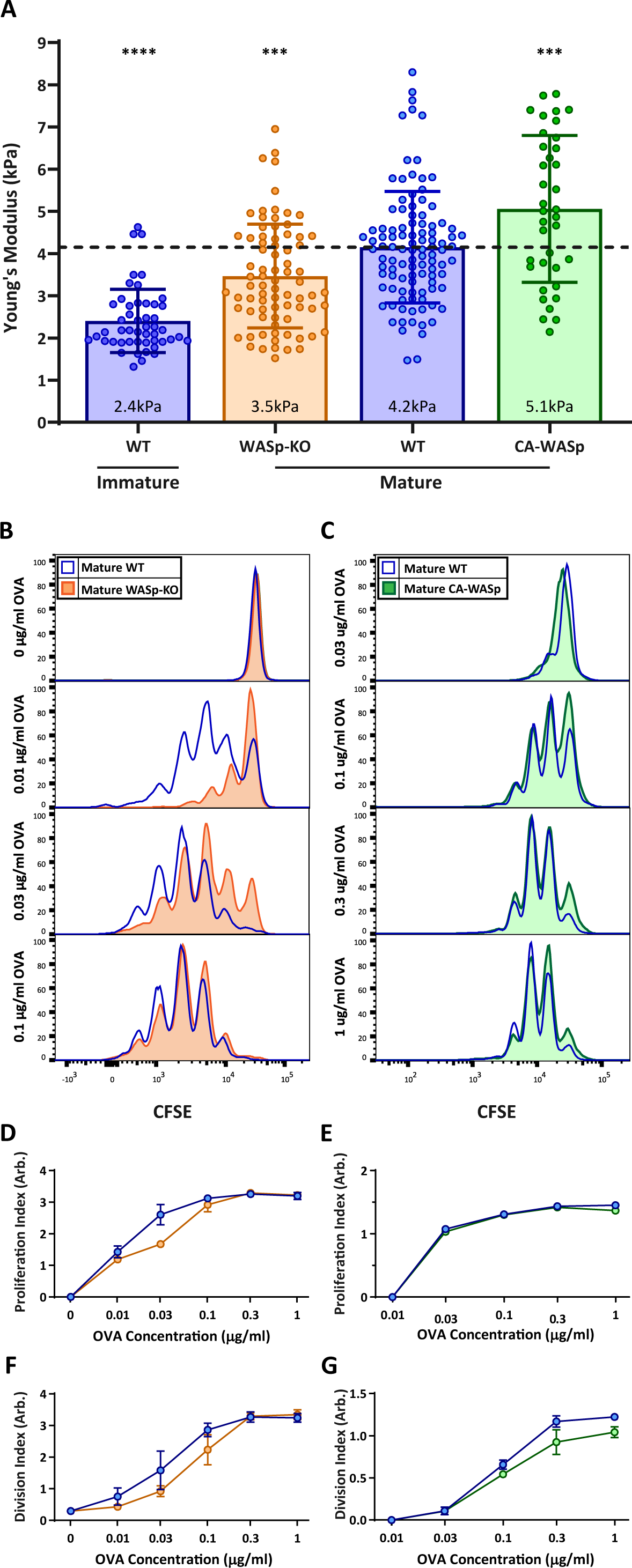
DC cortical stiffness acts as a costimulatory signal for T cell activation WT or WASp^-/-^ BMDCs, or WT BMDCs transduced with constituently active form of WASp (CA-WASp) were untreated or matured by treatment with LPS. (A) Cortical stiffness was measured by AFM micro-indentation. Each data point represents an average of two stiffness measurements at different locations around a single cell nucleus. Error bars denote standard deviation. ***p<0.001, ****p<0.0001 calculated by an unpaired one-way ANOVA, comparing mature WT with all other treatments, with post-hoc Tukey correction. (B,C) LPS-matured BMDCs were pulsed with a range of OVA_323-339_ peptide concentrations and co-cultured with *ex-vivo* OT-II CD4^+^ T cells for 72 hrs. Proliferation was measured by CFSE dilution. (D,E) Proliferation index pooled from two independent experiments. (F-G) Division index values pooled from two independent experiments. Error bars represent StDev.

## DISCUSSION

Recent work from several labs clearly shows that T cell activation involves mechanical cues. We have previously shown that the DC cytoskeleton constrains the mobility of stimulatory ligands on the DC surface, enhancing T cell activation by opposing the forces exerted by the T cell on the corresponding receptors (Comrie et al., 2015). In the current study, we elucidate a second mechanism whereby the DC cytoskeleton enhances T cell activation. We show that actin remodeling during DC maturation increases the cortical stiffness of DCs by 2-3 fold, and that T cell activation is enhanced by increases in stiffness over the same range. Importantly, increased stiffness lowers the threshold dose of TCR ligand needed for T cell activation, as expected if substrate stiffness serves as a costimulatory signal. In keeping with this concept, CD4^+^ T cells showed more profound stiffness-sensitivity than CD8^+^ T cells, especially at early times in the activation process. Together, these results indicate that stiffening of the DC cortex during maturation provides biophysical cues that work together with canonical costimulatory cues to enhance T cell priming.

Modulation of actin architecture has long been appreciated as an essential feature of DC maturation. Changes in the DC actin cytoskeleton facilitate the transition from highly endocytic tissue-resident cells to migratory cells specialized for antigen presentation (Burns et al., 2004; Burns et al., 2001). Our findings reveal a new facet of this process. We show that immature DCs are very soft, and that upon maturation, their cortical stiffness is increased by 2-3 fold. This is true for cultured BMDCs treated with LPS *in vitro*, as well and splenic DCs harvested from LPS-treated mice. A similar trend was reported by Bufi et al for human monocyte-derived DCs (Bufi et al., 2015), although that study reported lower absolute Young’s modulus values. While we used AFM indentation, Bufi et al. used microplate rheology. Since different methods for measuring cell mechanical properties produce absolute Young’s modulus values that can vary by as much as 100 fold (Wu et al., 2018), it seems likely that the apparent discrepancy in absolute values stems from technical differences between the two studies. Nevertheless, it is clear from both studies that the stiffness of the DC cortex is modulated during maturation.

The observed increase in stiffness depends on changes in actin architecture; whereas depolymerization of actin filaments has no effect on the stiffness of immature DCs, the increase associated with maturation depends on intact filaments, and is sensitive to inhibitors of actin polymerizing molecules. While it remains to be determined exactly which actin regulatory pathways control cortical stiffness in mature DCs, our data show that both Arp2/3 complex and formins are involved. Moreover, we find that DCs lacking the Arp2/3 activator WASp are abnormally soft. In keeping with these findings, DC maturation is known to induce changes in the activation state and localization of Rho family GTPases, especially Cdc42, a molecule that can activate both WASp and formins (Garrett et al., 2000; Vargas et al., 2016; West et al., 2000). Since the overall levels of active Cdc42 are diminished during DC activation, it seems likely that the observed increase in cortical stiffness results from redistribution of the active pool.

We show that DC cortical stiffness is a cell-intrinsic property that is unaffected by substrate stiffness. In this respect, DCs are different from other cell types that adapt their stiffness to differences in substrate compliance (Byfield et al., 2009; Tee et al., 2011). The ability of DCs to maintain constant stiffness despite changing environmental cues is reminiscent of previous work showing that DCs rapidly change their method of locomotion in order to maintain consistent migration speed and shape while crossing over different surfaces (Renkawitz et al., 2009). This behavior has been proposed to allow DCs to pass through tissues with widely different mechanical properties. In the same way, we propose that the ability of DCs to regulate cortical stiffness as a function of maturation state in spite of environmental cues reflects the importance of this property for priming an appropriate T cell response.

A central finding of this paper is that changes in DC stiffness serves as a costimulatory signal for T cell priming. By using a matrix of different hydrogels spanning the biologically relevant range defined for immature and mature DCs (2 – 8 kPa), coated with increasing pMHC concentrations, we found that stimulatory substrates with lower stiffness required higher concentrations of pMHC to achieve T cell activation. Similarly, when compared to WT DCs, softer WASp knockout DCs required higher concentrations of OVA peptide to induce the same level of proliferation.

Our results indicate that increases in cortical stiffness, together with diminished ligand mobility (Comrie et al., 2015), represent biophysical cues that are modulated in parallel with upregulation of costimulatory ligands and cytokines as a fundamental part of DC maturation. When interacting T cells engage pMHC complexes and costimulatory ligands on the DC surface, they integrate this biophysical input along with other canonical costimulatory signals.

In addition to lowering the antigenic threshold for T cell activation, changes in stiffness may present a new signaling mechanism by which DCs control T cell fate and differentiation. Bufi et al. showed previously that human monocyte-derived DCs responding to different maturation signals vary in their stiffness (Bufi et al., 2015). Interestingly, they found that treatment with the tolerizing cytokines TNFα and prostaglandin E2 results in DCs that are even softer than immature cells. Tolerogenic DCs exhibiting partially immature phenotypes have been shown to induce differentiation of regulatory T cells (Doan, McNally, Thomas, & Steptoe, 2009; Gleisner, Rosemblatt, Fierro, & Bono, 2011; Gordon, Ma, Churchman, Gordon, & Dawicki, 2014). This effect is usually attributed to low expression of T cell ligands or cytokines, but based on our data, we propose that biophysical properties of the DC cortex also play a role. Going forwards, it will be important to ask how DC stiffness is modulated in response to different environmental cues, and whether this further shapes T cell responses.

While we demonstrate T cell stiffness responses on soft surfaces emulating DCs, others have reported T cell stiffness responses on very stiff surfaces (Judokusumo et al., 2012; O’Connor et al., 2012). We found that very stiff substrates (25kPa hydrogels and plastic surfaces in the GPa range) elicit strong responses. This was true for proliferation, IL-2 secretion and degranulation. Similarly, recent analysis of human CD4^+^ effector T cells shows that re-stimulation on soft surfaces induces upregulation of genes related to cytokine signaling and proliferation, while restimulation on very stiff surfaces (100 kPa) triggers expression of an additional genetic program that includes metabolic proteins related to glycolysis and respiratory electron transport (Saitakis et al., 2017). The physiological relevance of these augmented responses is unclear, since T cells probably never encounter such stiff stimulatory surfaces *in vivo*. Nonetheless, such findings raise important questions about traditional *in vitro* assays of T cell function, which often utilize glass or plastic stimulatory surfaces.

The observation that T cells respond to APC stiffness is best understood in the context of evidence that T cells exert force on an interacting APC through the TCR complex (Bashour et al., 2014; Blumenthal & Burkhardt, 2020; Hui et al., 2015; Husson et al., 2011; Li et al., 2010; Sawicka et al., 2017), with the amount of force corresponding to APC stiffness (Husson et al., 2011; Sawicka et al., 2017). Apart from being a requirement for activation (Li et al., 2010; Pryshchep et al., 2014), force transduction has been shown to promote peptide discrimination by influencing bond lifetimes (Das et al., 2015; Liu et al., 2014). Importantly, it appears that the TCR’s ability to sense stiffness is closely related to its ability to transduce force-dependent signals during T cell-APC interaction. Indeed, there is evidence that signaling downstream of TCR engagement is increased on stiffer substrates (Judokusumo et al., 2012) and that the location of early tyrosine phosphorylation events corresponds to sites of maximum traction force (Bashour et al., 2014). We propose that stiffer substrates allow T cells to exert more force through TCR interactions, and consequently induce more effective signaling. This accounts for the co-stimulatory property of substrate stiffness on T cell activation.

The mechanism by which force application on the TCR is translated into biochemical signals remains controversial. Nevertheless, there is evidence to suggest that force applied on the TCR complex induces conformational changes within TCRαβ that exposes ITAM sites on the CD3 and TCRζ chains for phosphorylation and downstream signaling ^57,58.^ Importantly, conformational changes are mainly attributed to extension of the CβFG loop region within TCRβ (Das et al., 2015), which serves as a lever to push down on the CD3 complex (Sun, Kim, Wagner, & Reinherz, 2001), exposing ITAM sites (Xu et al., 2008). In support of this idea, we found that the way in which the TCR is engaged influences T cell stiffness sensing. Within the biologically relevant stiffness range (2-8 kPa), T cells were activated only when TCRαβ was engaged directly; indirect engagement through anti-CD3 resulted in almost no response. We postulate that direct TCRαβ engagement leads to conformational changes that are transmitted appropriately for efficient initiation of downstream signaling, whereas engagement of CD3 induces smaller changes and more limiting downstream signaling. This effect may be most evident on soft substrates, because force-dependent signaling is limiting in this setting. We note that on these soft substrates, pMHC induced stronger T cell activation than anti-TCRβ. This may reflect the involvement of CD4 in the former, which leads to more efficient recruitment of Lck to the TCR complex.

Although our focus here is on the role of stiffness sensing in priming of naïve T cells, we find that effector CD8^+^ T cells also exhibit stiffness-dependent degranulation responses. Similarly, Saitakis et al. have shown that restimulating CD4^+^ effector T cells on surfaces of different stiffness induces differential gene expression and cytokine production (Saitakis et al., 2017). Since DCs increase their cortical stiffness during maturation, a stiffness dependent mechanism for naïve T cell priming makes biological sense. Effector T cells, however, interact with a variety of APCs. In particular, cytotoxic CD8^+^ T cells are expected to kill any infected cell throughout the body with no stiffness bias. The physiological significance of stiffness sensitivity for effector T cells remains unclear. One intriguing possibility is that mechanosensing is an obligate component of the feedback loop that underlies force-dependent TCR triggering.

The IS is often described as a platform for information exchange between the T cell and APC. Together with our recent work on ligand mobility, the findings presented here indicate that the mechanical properties of the APC side of the IS influence T cell priming, likely because they augment force-dependent conformational changes in TCRs, integrins, and potentially other molecules. Going forward, it will be important to determine how these properties are modulated during DC maturation, and whether there are also local changes induced by signaling events taking place at the IS. In addition, it will be important to tease apart the molecular events through which T cells sense and respond to these mechanical cues.

### MATERIALS AND METHODS

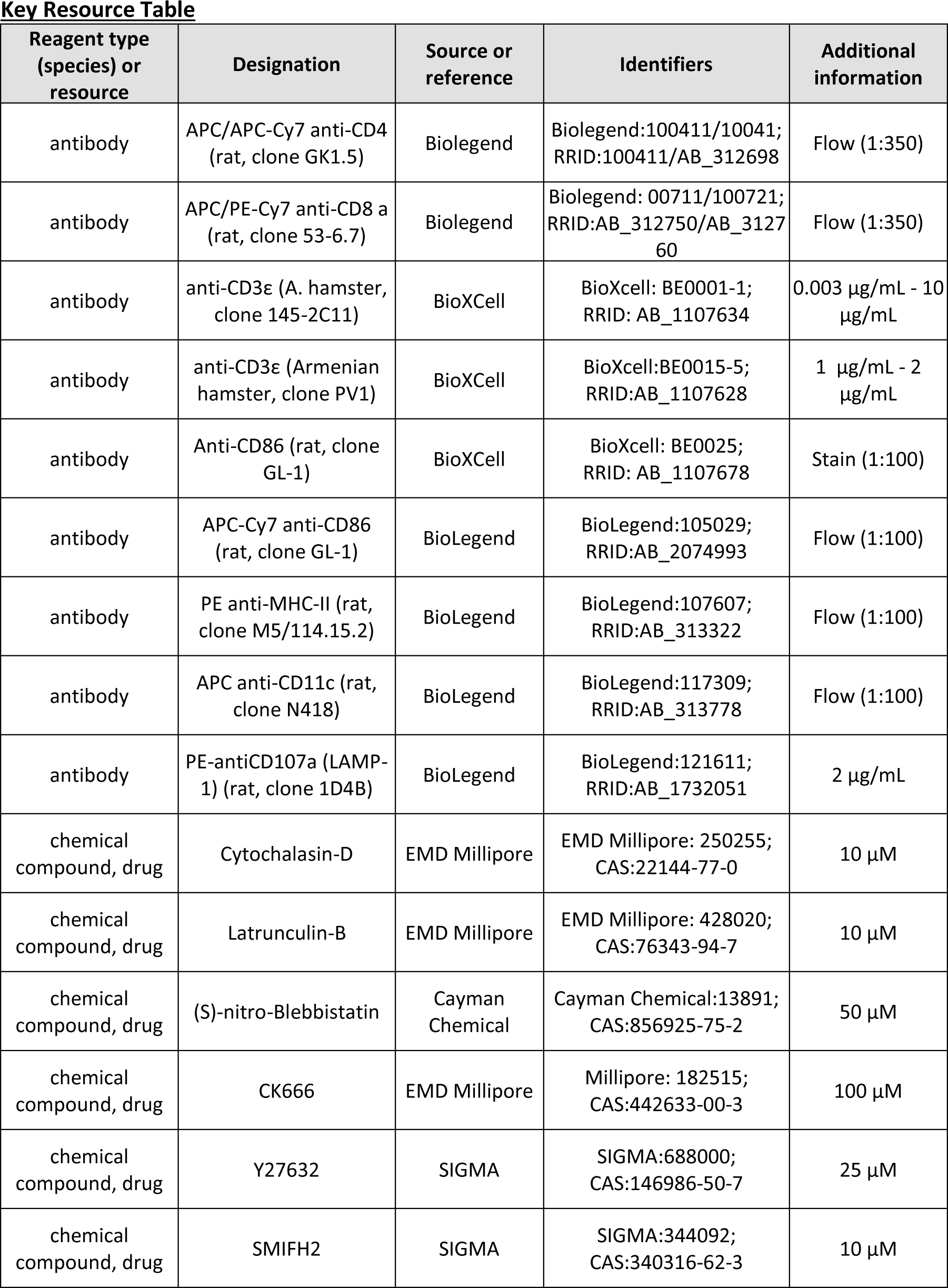
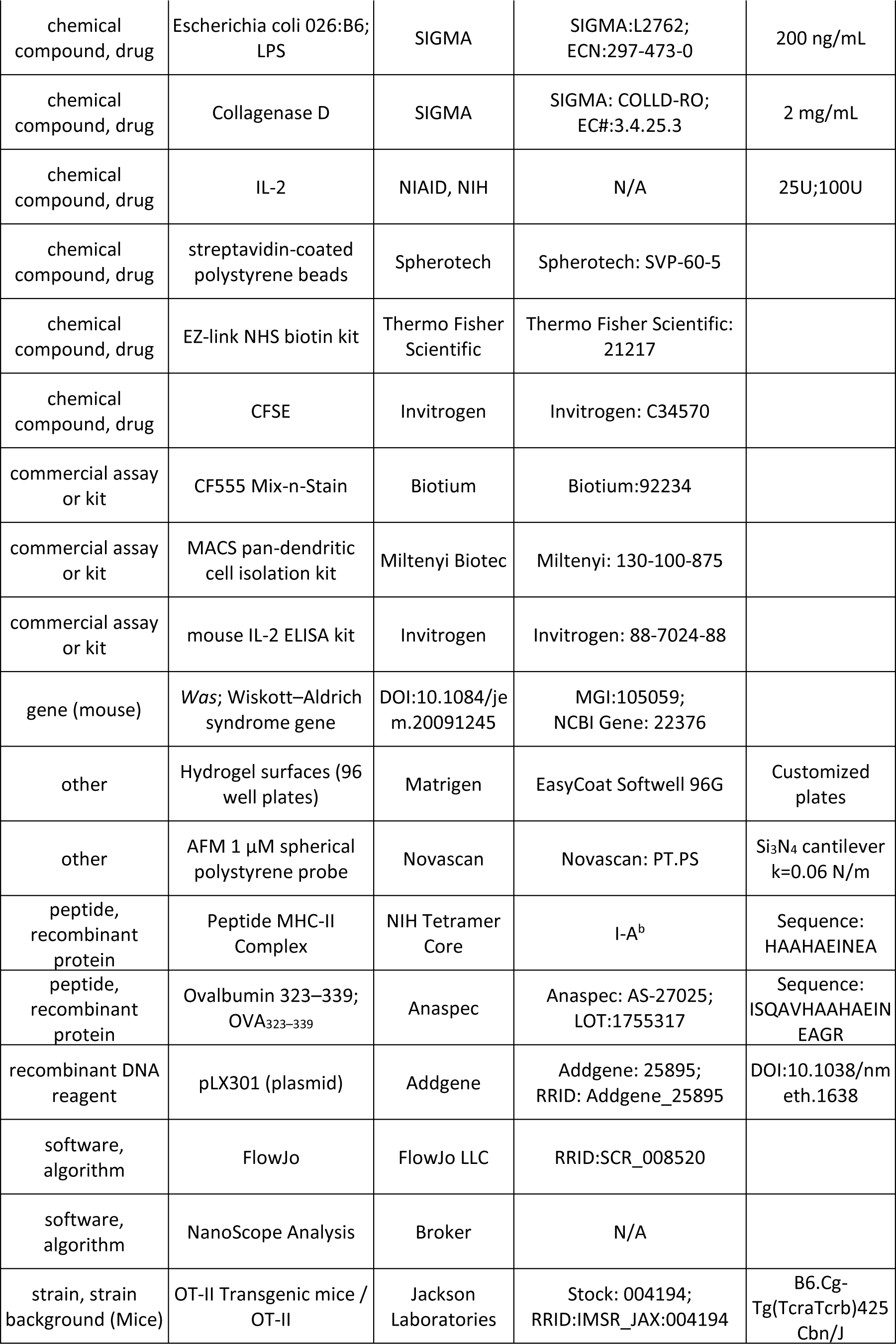
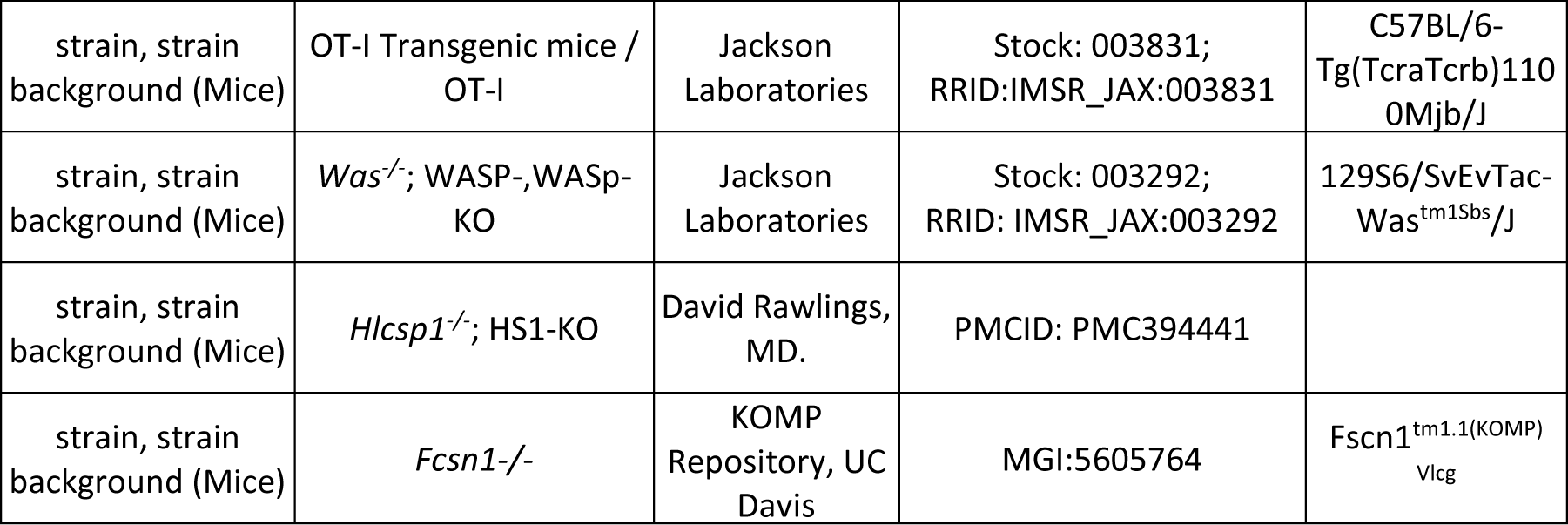

### Inhibitors and antibodies

Cytochalasin-D and Latrunculin-B were from EMD Millipore, (S)-nitro-Blebbistatin was from Cayman Chemical, CK666 was from Calbiochem, and Y27632 and SMIFH2 were from Sigma-Aldrich. <u>Flow cytometry antibodies:</u> rat anti-CD4 APC/APC-Cy7 (clone RM4-5), rat anti-CD8a APC/PE-Cy7 (clone 53-6.7), Armenian hamster anti-CD69 APC (clone H1.2F3) and rat anti-CD25 PE (clone PC61) were all from BioLegend. <u>Surface coating antibodies</u>: Armenian hamster anti-CD3ε (clone 2C11), and Armenian hamster anti-CD28 (Clone PV1) were from BioXCell. Biotinylated Armenian hamster anti-CD3ε (clone 2C11) was from Invitrogen, and biotinylated mouse anti-TCRVβ5.1/5.2 (clone MR9-4) was from BD Bioscience. <u>Dendritic cell staining:</u> anti-CD86 CF555 was made by conjugating purified rat anti-CD86 (BioXCell) with CF555 conjugated dye from Biotium as per the manufacturer’s protocol.

### Mice

All mice were originally obtained from The Jackson Laboratory and housed in the Children’s Hospital of Philadelphia animal facility, according to guidelines put forth by the Institutional Animal Care and Use Committee. C57BL/6 mice (WT) were purchased from Jackson Labs. HS1-KO mice on the C57BL/6 background have been previously described(Taniuchi et al., 1995) and were a kind gift from doctor David Rawlings at the University of Washington. *Was*^-/-^ mice were purchased from Jackson labs (Snapper et al., 1998) and fully backcrossed to a C57BL/6 background. All mouse strains were used as a source of bone marrow from which to generate BMDCs. Mice bearing a gene trap mutation in the Fscn1 gene (Fscn1^tm1(KOMP)Vlcg^), which abrogates expression of the protein Fascin 1, were generated by the KOMP Repository at UC Davis, using C57BL/6 embryonic stem cells generated by the Texas A&M Institute for Genomic Medicine. Because these mice proved to have an embryonic lethal phenotype, fetal liver chimeras were used as a source of bone marrow precursors. Heterozygous mating was performed, and fetal livers were collected after 15 days of gestation and processed into a single-cell suspension by mashing through a 35-μm filter. Embryos were genotyped at the time of harvest. Cells were resuspended in freezing media (90% FCS, 10% DMSO) and kept at −80 °C until used. Thawed cells were washed, counted, resuspended in sterile PBS and injected i.v. into sub-lethally irradiated 6-week-old C57BL/6 recipients, 1 × 10^6^ cells per mouse. Chimeras were used as a source for fascin KO bone marrow ∼6 weeks after transfer. OT-I T cells were prepared from heterozygous OT-I TCR Tg mice, which express a TCR specific for ovalbumin 257-264 (amino acid sequence SIINFEKL) presented on H-2K^b^ (Hogquist et al., 1994). OT-II T cells were prepared from heterozygous OT-II TCR Tg mice, which express a TCR specific for ovalbumin 323–339 (amino acid sequence ISQAVHAAHAEINEAGR) presented on I-A^b^ (Barnden, Allison, Heath, & Carbone, 1998).

### Cell culture

Unless otherwise specified, all tissue culture reagents were from Invitrogen/Life Technologies. GM-CSF was produced from the B78H1/GMCSF.1 cell line (Levitsky et al., 1996). HEK-293T cells (ATCC) were cultured in DMEM supplemented with 10% FBS, 25mM Hepes, penicillin/streptomycin, GlutaMAX, and non-essential amino acids.

Generation of bone marrow derived dendritic cells (BMDCs) was similar to (Inaba et al., 1992). Briefly, mouse long bones were flushed with cold PBS, the resulting cell solution was passed through a 40 µm strainer, and red blood cells were lysed by ACK lysis. Cells were washed once with RPMI-1640 and then either frozen for later use in RPMI-1640 containing 20% FBS, 10% DMSO, or plated in 10 cm bacterial plates in BMDC culture media (RPMI-1640, 10% FBS, penicillin/streptomycin, GlutaMax and 1% GM-CSF supernatant). On day 3 of culture, dishes were supplemented with 10 ml of BMDC culture media. On Day 6, 10 ml of media were replaced, by carefully collecting media from the top of the dish and slowly adding fresh media. BMDC differentiation was verified by flow cytometry, showing 80-90% CD11c positive cells. BMDC maturation was induced on days 7 or 8; immature BMDCs were harvested and re-plated in a 6cm tissue culture dish in 5 ml of BMDC media supplemented with 200 ng/ml LPS (*Escherichia coli* 026:B6; Sigma-Aldrich) for at least 24h. Maturation was verified by flow cytometry, with mature BMDCs defined as Live/CD11c^+^/CD86^high^/MHC-II^High^ cells. To generate splenic DCs, spleens from C57BL/6 mice were cut to smaller pieces and digested with Collagenase D (2 mg/ml, Sigma) for 30 min at 37°C, 5%CO_2_. Cells were washed and labeled for separation by negative selection using a MACS pan-dendritic cell isolation kit (Miltenyi Biotec).

Primary mouse T cells were purified from lymph nodes and spleens using MACS negative selection T cell isolation kits (Miltenyi Biotec). In the case of CD4^+^ T cells, ex vivo cells were used, as isolation yielded high fractions of naïve cells. In the case of CD8^+^ T cells, naïve T cells were isolated, as ∼45% of T cells isolated from OT-I mice showed some level of activation. To generate cytotoxic CD8^+^ T cells (CTLs), purified murine CD8^+^ cells were activated on 24-well plates coated with anti-CD3ε and anti-CD28 (2C11 and PV1, 10μg/ml and 2 µg/ml respectively) at 1×10^6^ cells per well. After 24h, cells were removed from activation and mixed at a 1:1 volume ratio with complete T cell media (DMEM supplemented with penicillin/streptomycin, 10% FBS, 55μM β-mercaptoethanol GlutaMAX, and non-essential amino acids), containing recombinant IL-2 (obtained through the NIH AIDS Reagent Program, Division of AIDS, NIAID, NIH from Dr. Maurice Gately, Hoffmann – La Roche Inc (Lahm & Stein, 1985)), to give a final IL-2 concentration of 100 units/ml. Cells were cultured at 37°C and 10% CO_2_, and passaged as needed to be kept at 0.8×10^6^ cells/ml for 7 more days. CTLs were used at days 8 or 9 after activation.

### Plasmid construction, viral production, and transduction of DCs

A constitutively active form of WASp (CA-WASp) was engineered by subcloning WASp cDNA into a pLX301 vector (Addgene), introducing an I294T point mutation (Westerberg et al., 2010) by site-directed mutagenesis, and confirming by sequencing. To generate recombinant lentivirus, HEK-293T cells were co-transfected using the calcium phosphate method with psPAX2 and pMD2.G, together with the DNA of interest in pLX301. For transduction, BMDCs were plated in untreated 6 well plates at 2×10^6^ cells/well in 3 ml of BMDC media. BMDC transduction was carried out on day 2 of culture; lentiviral supernatants were harvested from HEK-293T cells 40hr post transfection, supplemented with 8 µg/ml Polybrene (Sigma-Aldrich), and used immediately to transduce BMDCs by spin-infection at 1000xG, 37°C for 2hr. After resting the cells for 30 min at 37°C, 5% CO_2_, lentivirus-containing media was replaced with normal BMDC culture media. On day 5 of culture, puromycin (Sigma-Aldrich) was added to a final concentration of 2 μg/ml to allow selection of transduced BMDCs. Maturation of transduced cells was induced on day 8 by adding 200 ng/ml of LPS in puromycin-free media.

### Flow Cytometry

All cells were stained with Live/Dead aqua (ThermoFisher) following labeling with appropriate antibodies in FACS buffer (PBS, 5% FBS, 0.02% NaN_3_, and 1 mM EDTA). Flow cytometry was performed using either the Cytoflex LX or CytoFlex S cytometer (Beckman Coulter) and analyzed using FlowJo software (FlowJo LLC). T cells were gated based on size, live cells, and expression of CD4 or CD8 (depending on the experiment). DCs were incubated for 10 min on ice with the Fc blocking antibody 2.4G2 prior to staining. DCs were gated based on size, live cells, and CD11c expression. Mature DCs were further gated based on high expression of MHC-II, CD86 or CD80.

### T cell activation on stimulatory gel surfaces

96 well plates coated with polyacrylamide hydrogels spanning a stiffness range of 2 – 25 kPa were obtained from Matrigen. Hydrogel stiffness was verified by AFM at different locations around the hydrogel surface (Figure 3 - Figure Supplement 1). Surfaces were first coated with 10 µg/ml of NutrAvidin (ThermoFisher) and 2 µg/ml anti-CD28 (clone PV1) overnight at 4°C. Primary amines in the NutrAvidin form covalent bonds with quinone functional groups within the hydrogels. The gel pore size is on the order of tens of nanometers, such that cells can only interact with ligands bound on the gel surface. Surfaces were then washed twice with 200 µL of PBS and coated with varying concentrations of biotinylated pMHC monomers (NIH tetramer core facility) for 2 hours at 37°C. Where antibodies were used for coating, surfaces were coated with varying concentrations of either biotinylated anti-CD3ε (clone 2C11), or biotinylated anti-TCRvβ5.1/5.2 (clone MR9-4). Surfaces were than washed 2 times with 200 µL of PBS, and blocked for 10 min with T cell media containing 10% FBS prior to addition of 2.0×10^5^ cells/well. Control studies showed that stimulatory ligands bound slightly less well to stiffer hydrogel surfaces (Figure 3 - Figure Supplement 2). Stiffer surfaces show the same or higher activating properties across all assays, ruling out the possibility that differences in T cell activation are due to differential ligand binding. Importantly, initial experiments included a 1 kPa hydrogel surface that yielded no response across all assays. Therefore, data from this condition is not shown and was not included in repeated experiments. For experiments where exogenous IL-2 was added, media was supplemented with IL-2 to a final concentration of 25 U/ml. To measure IL-2 secretion, supernatants were harvested 22-24 hours post stimulation, and IL-2 concentration was measured using a mouse IL-2 ELISA kit (Invitrogen). For early activation marker expression assays, cells were plated immediately after isolation, and harvested 22-24 hours post stimulation for flow cytometry analysis. For CFSE dilution assays, purified cells were washed once with PBS and stained for 3 min with 2.5 µM CFSE (ThermoFisher). After quenching the excess CFSE by addition of 1 ml FBS for 30 seconds, cells were washed and plated. Cells were harvested 68-72 hours post stimulation for flow cytometry analysis. For ligand comparison assays, surfaces were first coated with 10 µg/ml of NutrAvidin (ThermoFisher) and 2 µg/ml anti-CD28 (PV1) overnight at 4°C. Surfaces were then washed twice with 200 µL of PBS and coated with varying concentrations of biotinylated ligands (anti-TCRVβ5.1/5.2, anti-CD3ε, or pMHC-II monomers) for 2 hours at 37°C.

### Cytotoxic T cell degranulation assays

Assays were conducted on day 8 or 9 of culture. 2×10^5^ CTLs were plated onto surfaces coated with various concentrations of pMHC-I in the presence of 2 µg/ml PE-conjugated anti-CD107a. After 3 hours of re-stimulation, CD107a labeling was quantified by flow cytometry analysis. Cells were gated based on size, live cells, and expression of CD8^+^. CD107a mean fluorescence intensity (MFI) was extracted using FlowJo.

### T cell priming assays

Priming assays were carried out in round bottom 96 well plates. 5×10^4^ LPS-matured BMDCs were plated in each well and pulsed with OVA_323–339_ peptide at various concentrations (0.1 – 1 µg/ml). 1.5×10^5^ CFSE stained, OT-II CD4^+^ T cells were added to each well and incubated for 68-72 hours. Cells were then harvested and analyzed using flow cytometry.

### Atomic force microscopy (AFM)

All experiments were carried out at room temperature using a Bruker Bioscope Catalyst AFM mounted on a Nikon TE200 inverted microscope. Micro-indentation measurements were made with a spherical tip from Novascan. The tip was comprised of a 1 µm silicon dioxide particle mounted on a silicon nitride cantilever with a nominal spring constant of 0.06 N/m; each cantilever was calibrated using the thermal fluctuation method. The AFM was operated in fluid contact mode, with 2 Hz acquisition. Total vertical cantilever displacement was set to 5 µm, producing a maximal approach/retraction speed of ∼20 µm/sec. Maximal deflection (Trigger threshold) was adjusted for each cantilever to apply a maximal force of 6 nN on the measured cell (e.g. for a 0.06 N/m cantilever, the trigger threshold was set to 100 nm). The actual indentation depth was ∼1.5 µm depending on the measured cell stiffness (Figure 1 – Supplement Figure 2). Analysis of force-distance curves was carried out using the Nanoscope Analysis software (Bruker). The Young’s modulus was extracted using the Hertzian model for spherical tips with a contact point-based fitting on the extend curve data. For each individual cell, two separate measurements were conducted at different locations near, but not directly over the nucleus. The reported cell stiffness value represents the average between these independent measurements. Note that when measurements of cortical stiffness were made over the nucleus, no significant differences in Young’s modulus values were found (not shown). To measure BMDC stiffness, 1×10^5^ cells (untreated or LPS matured) were seeded onto Poly L-lysine coated coverslips and allowed to spread for 4 hours at 37°C, 5% CO_2_. Prior to data acquisition, cells were incubated for 10 min with the Fc blocking antibody 2.4G2, washed and stained for CD86 for 20 min, then washed and mounted on the AFM. All antibody incubations and data acquisition steps were performed in L-15 media (Gibco) supplemented with 2mg/ml glucose. For treated cell measurements, drugs [Latrunculin-B (10 µM), Cytochalasin-D (10 µM), s-nitro-Blebbistatin (50 µM), Y27632 (25 µM), CK666 (100 µM), or SMIFH2 (10 µM)] were pre-incubated with the cells at 37°C, 5% CO_2_ for 30 min prior to Fc blocking and maintained in the cultures throughout staining and data acquisition.

### Statistical Methods

All datasets were subjected to outlier analysis prior to execution of statistical testing. Outliers were defined as data points with values outside the range of mean +/-2.5xStDev, and were deleted from the dataset. Testing for a statistically significant difference between experimental groups was done using an unpaired one-way ANOVA test with a post-hoc Tukey correction for multiple comparisons.

Throughout the paper, data shown represents biological, and not technical, replicates. For BMDC assays, a single experiment constitutes measurement of multiple cells from a fresh DC culture, starting from frozen or freshly harvested bone marrow. For splenic DCs, a single experiment constitutes measurement of multiple cells freshly purified from the spleen of a single mouse. In each experiment, WT or untreated cells were measured side by side with treated cells as a standard control. For T cell assays, a single experiment constitutes cells freshly purified from spleen and lymph nodes of a single animal. All CFSE dilution assays were executed in technical duplicates, although a single data set is presented. When needed, figure legends describe the quantity of technical repeats used in an experiment.

## ACKNOWLEDGEMENTS

The authors thank Florin Tuluc and Jennifer Murray from the Children’s Hospital of Philadelphia Flow Cytometry core. We thank the biomechanics core of the Institute of Translational Medicine and Therapeutics (ITMAT) at the University of Pennsylvania for use of the Atomic Force Microscope and the NIH tetramer core facility for provision of MHC-peptide complexes. We thank Dr. Shuixing Li and Dr. Nathan Roy for expert technical assistance, Dr. Nathan Roy and Mr. Tanner Robertson for critical reading of the manuscript, and members of the Burkhardt laboratory for many helpful discussions. This work was supported by NIH grants R01 GM104867 and R21 AI32828 to JKB.

## ETHICS STATMENT

All studies, breeding and maintenance of animals was performed under Animal Care and Use Protocol #667, as approved by The Children’s Hospital of Philadelphia Institutional Animal Care and Use Committee.

---

**Figure 1 – Figure Supplement 1. Validation of hydrogel compliances.** Direct AFM measurements of hydrogel compliance, validating the stiffness of hydrogels used for measuring the effects of substrate compliance on DC stiffness

**Figure 1 – Figure Supplement 2. Indentation length in AFM measurements of DC cortical stiffness.** Indentation length was defined as the distance between initial deflection to maximal cantilever displacement. (A) an example of a force curve and the extraction of the *de-facto* indentation length. (B) quantification of indentation length for immature and mature DCs.

**Figure 3 – Figure Supplement 1. Validation of hydrogel compliances.** Direct AFM measurements of hydrogel compliance, validating the stiffness of hydrogel surfaces used for activation of T-cells.

**Figure 3 - Figure Supplement 2. Binding of stimulatory antibodies to various hydrogel surfaces** Hydrogel surfaces were coated with 10 ug/ml of DyLight800 conjugated NutrAvidin (ThermoFisher) for 2.5 hours, than washed 3 times with 200 µL of PBS, with 5 min incubations. Imaging was done on a LI-COR Odyssey reader. (A) Fluorescence image of coated and uncoated hydrogel plates (B) Quantification of mean fluorescent intensity for the center of each well. Each data point represents mean +/-StDev pooled from two independent hydrogel wells.

