## Supplementary material for "T cell priming is enhanced by maturation-dependent stiffening of the dendritic cell cortex": Figure 1 - Supplemental Figure 1

### FIgure 1 - Figure Supplement 1

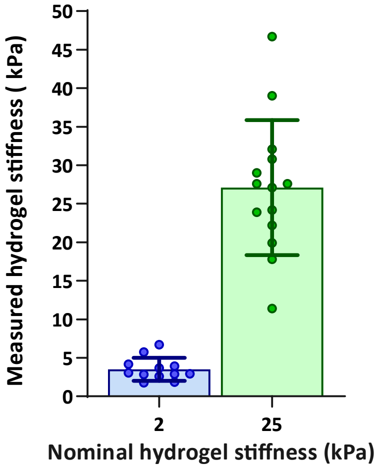
