## Supplementary material for "T cell priming is enhanced by maturation-dependent stiffening of the dendritic cell cortex": Figure 3 - Supplemental Figure 1

### Figure 3 - Figure Supplement 1

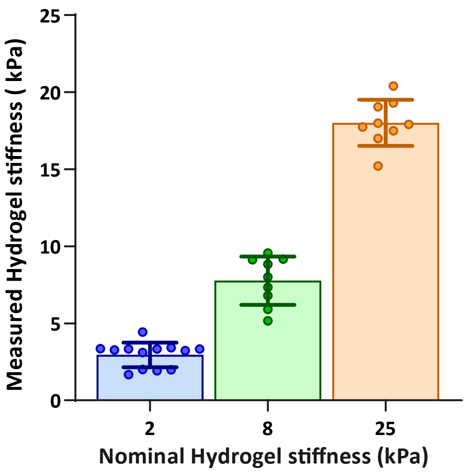
