## Supplementary figures and images for "T cell priming is enhanced by maturation-dependent stiffening of the dendritic cell cortex"

### Figure 3 - Supplemental Figure 2

#### Figure 3 - Figure Supplement 1

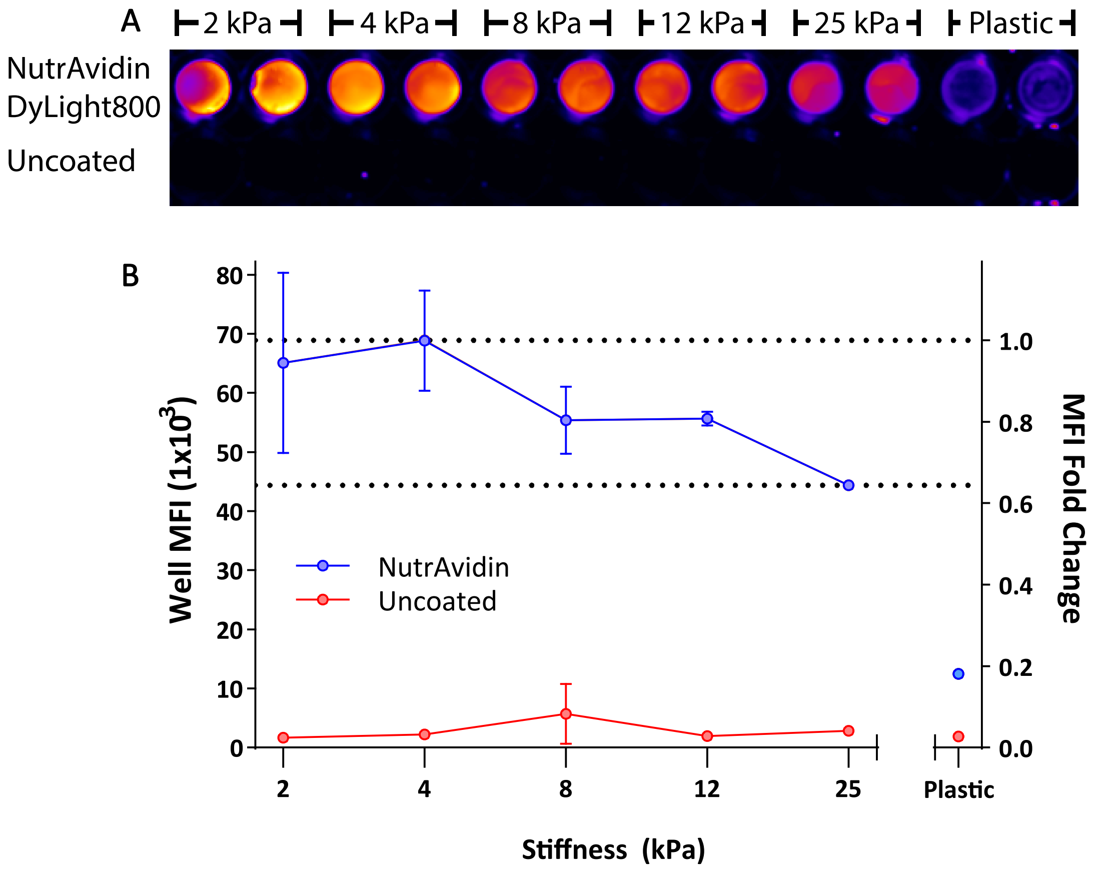
